# Learning of visual sequences by neurons in the human hippocampus

**DOI:** 10.1101/2025.03.04.641300

**Authors:** Tibin John, Yajun Zhou, Ayman Aljishi, Bastian Rieck, Nicholas B. Turk-Browne, Eyiyemisi C. Damisah

## Abstract

Time is a critical component of memory and yet how the hippocampus incorporates temporal information with the sensory contents of memories remains unclear. We hypothesized that the hippocampus can learn arbitrary sequences through rapid changes in the tuning and population geometry of individual neurons to mirror the sequence structure. We recorded single-unit activity from 134 neurons in the hippocampus of 17 patients with epilepsy while they viewed sequences of visual scenes that were presented repeatedly in the same order by looping from the end to start. As the sequence repeated, hippocampal neurons that were initially responsive to one scene in the sequence began responding to the other scenes proportional to their temporal proximity. The population vectors of spiking activity across recorded hippocampal neurons for each scene came to form a high-dimensional ring topology that encoded the circular structure of the sequence in a manner that preserved the serial order of the scenes. These effects were not observed when sequences were scrambled upon each repetition to destroy temporal structure, nor in brain regions outside of the hippocampus for both structured and random sequences. These findings suggest that temporal structure governs the representation of sensory stimuli in human hippocampal neurons.

## Introduction

Memory involves binding elements of an experience over space and time (1–5). Accordingly, neurons in the hippocampus — an essential brain region for memory — represent objects, concepts, people, and other sensory features (6); individual locations in an environment (7); specific moments in time during an event (8); and conjunctions of these features organized into hierarchical, contextual representations (9–12). Although the coding of spatial information in hip-pocampal neurons has been characterized extensively over the past half century, their temporal coding has only been studied more recently, especially in humans. With single-unit recordings from the hippocampus of patients with epilepsy, these pioneering studies have revealed neurons that code for moments in time during an event (13) and serial positions (14) in a sequence.

Here we investigate how the human hippocampus combines its sensitivity to time and sensory features in order to learn repeating visual sequences. We hypothesize that individual hip-pocampal neurons represent the temporal position of items in the sequence. These neurons initially respond to one item but begin responding to other items in the sequence across repetitions proportional to their temporal proximity. We further hypothesize that these changes result in a population code that represents the temporal geometry and serial order of the sequence. Vectors corresponding to the collective response of hippocampal neurons for each item are arranged topologically in a manner that preserves sequence distances.

This hypothesis is based on electrophysiological studies of associative learning in higher-order visual cortex of non-human primates and on fMRI studies of statistical learning in the human hippocampus. The primate studies began with the serendipitous discovery that repeated exposure to a visual sequence led neurons in inferior temporal and perirhinal cortices that had a preference for one item to respond to neighboring items in the sequence, with the response decreasing proportional to the temporal distance of the neighbor (15). These neurons also showed increased activity to an arbitrary item that was paired with a preferred item in a paired associate task (16) or across saccadic eye movements (17).

The human studies involved exposing participants to a continuous visual sequence that contained hidden temporal regularities in terms of which items tended to follow each other (18). Items that were consistently paired over time came to evoke more similar patterns of high-resolution fMRI activity in the hippocampus after versus before sequence exposure, even when they were presented individually. The same logic was used to demonstrate that the hippocampus encodes more complex temporal regularities in the similarity of fMRI activity patterns, by exposing participants to visual sequences generated from walks on a community structure graph (19). A recent intracranial EEG study similarly found that the population activity of hippocampal neurons can represent graph structure and trajectories (20).

The current study builds on this prior work to characterize how single neurons in the human hippocampus change their tuning to encode repeating visual sequences, and how these changes manifest in population geometry. We establish the role of learning with a control condition that scrambles the sequence on each repetition and assess specificity to the hippocampus with recordings from other brain regions.

## Results

We recorded single-unit activity from the hippocampus of 17 patients with epilepsy who had implanted microwires (see Figure 1a, Table S1 for microwire locations and Table S2 for subject demographics). The participants completed a passive viewing task in which they watched a stream of photographs of natural scenes (Figure 1b) in blocks of two conditions. In the structured condition (Figure 1c), 10 scenes were presented in a fixed order that wrapped around at the end, resulting in a looping circular sequence that was repeated 15 times; the order was chosen randomly for the first time (and fixed for subsequent repetitions) and thus was arbitrary in terms of the perceptual and semantic relationships between scenes. In the random condition (Figure 1d), 10 other scenes were presented in a unique random order on each of the 15 repetitions. Thus, conditions were equated in the type, number, and frequency of items, varying only in whether the order of the scenes was predictable.

**Fig. 1.**
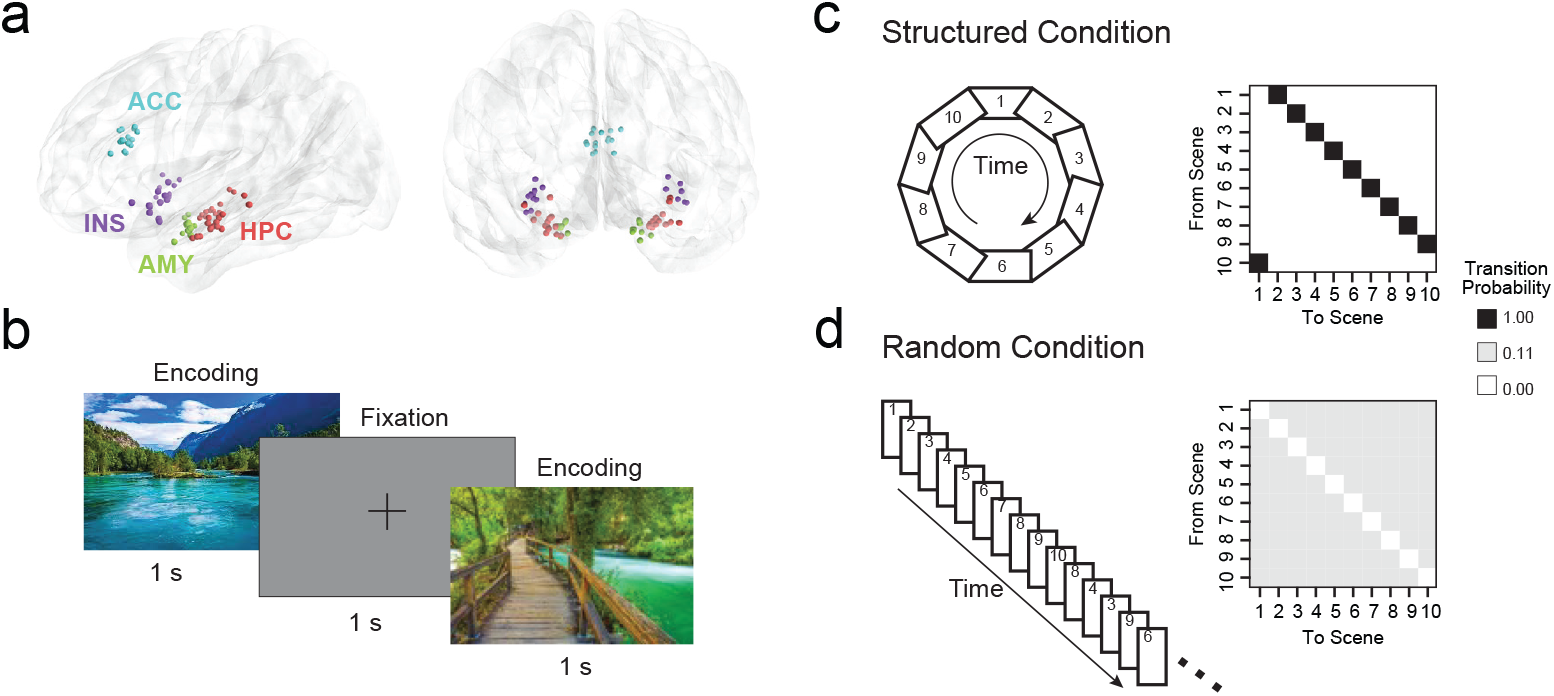
Recording locations and task design. (a) Brain reconstructions from 17 patients displaying the electrodes color-coded by brain region, including our primary region of interest, the hippocampus (HPC, red), and as control regions, the amygdala (AMY, green), insula (INS, purple), and anterior cingulate cortex (ACC, cyan). (b) Participants passively viewed streams of scene photographs, with each scene displayed for 1 s and separated by 1 s of fixation. (c) In the structured condition, the order of 10 scenes (numbered) was fixed and repeated 15 times in a loop, resulting in a reliable, circular sequence. This is reflected in the transition matrix as deterministic transitions between consecutive scenes. (d) In the random condition, the order of the scenes was randomized on each of the 15 repetitions eliminating sequential structure. Because back-to-back repetitions were prevented, the transition matrix is uniform over the nine remaining scenes.

### Initial responses of hippocampal neurons to scenes

We recorded a total of 444 neurons, including 134 from the hippocampus, 89 from the amygdala, 108 from the insula, and 113 from the anterior cingulate cortex (ACC). To assess sequence learning at the single-unit level, we first defined a putative reference scene for each neuron by identifying which of the 10 scenes in each condition evoked the highest mean firing rate across its first 3 presentations. A neuron was considered responsive to its reference scene only if the initial firing rate to that scene was significantly greater than: its neighboring scenes in the sequence, a fixation baseline period, and a null distribution generated by shuffling scene labels (see Methods). This selection procedure was intended to identify neurons whose tuning might undergo distance-dependent changes with sequence exposure, which was assessed by anchoring learning-related analyses of each neuron on its reference scene. In the hippocampus, 22 neurons in the structured condition and 31 neurons in the random condition met the criteria of being initially responsive to their reference scene. As further validation of this approach, we quantified scene selectivity across all trials using mutual information (MI) between firing rate and scene identity (21, 22). MI-based estimates of the proportion of visually selective neurons in the hippocampus were significantly greater than chance and broadly consistent with previous reports (Figure S3a). Importantly, MI computed from the first 3 presentations of a scene (used above to identify a neuron’s reference scene) was strongly correlated with the MI across all trials in the hippocampus, indicating that the initial responsiveness of neurons to a scene was relatively stable across the session (Figure S3b).

Because not all neurons met the initial responsiveness criteria above in both conditions, comparisons between structured and random conditions were made across partially overlapping sets of neurons rather than within individual units. That is, for single-unit analyses, each responsive neuron contributed one data point for each condition in which it was recorded. Note that the population-level analyses below included all hippocampal neurons regardless of criteria for selectivity.

### Changes in single-unit activity with sequence learning

Having identified hippocampal neurons with a well-defined reference scene, we next asked how repeated exposure to a visual sequence containing that scene would shape responses to other scenes in the sequence. To capture learning, we compared neuronal responses early in sequence exposure (first 3 trials of 15) versus late in exposure (last 3 trials).

We predicted that, in the structured but not random condition, the response of hippocampal neurons to scenes would increase from early to late exposure inversely proportional to their sequential distance from the neuron’s reference scene.

The heatmap of baseline-subtracted firing rates for an example neuron across trials and scene distances illustrates progressive modulation of responses with sequence learning (Figure 2a). In the structured condition, neuronal responses in the hippocampus became graded by distance from the reference scene, with systematic changes in the difference of late minus early firing rates that decreased monotonically with absolute scene distance (Figure 2b). The increase in firing rate at scene distance 1 was significantly stronger than in the random condition (Mann–Whitney U test, *p* = 0.033). To quantify the trial-by-trial evolution of this distance dependence, we computed, for each neuron and each trial, the Spearman correlation between firing rate and distance relative to the reference scene (Figure 2c, left). We then summarized the learning-related change by fitting a linear slope across trials to these rate–distance correlations (Figure 2c, right). In the structured but not random condition, hippocampal neurons showed progressively stronger distance-dependence over trials, as reflected by a systematic drift toward more negative correlations (Figure 2d, left). Across hippocampal neurons (Figure 2d, right), the change across trials was highly significant in the structured condition (Wilcoxon signed-rank test against zero slope, *p <* 0.0001) but not the random condition (*p* = 0.555), and was significantly stronger in structured than random conditions (Mann–Whitney U test, *p* = 0.022). Other recorded regions including the amygdala, insula, and ACC did not show significant distance-dependent modulation or a systematic rate–distance correlation progression in either condition (Figure S4). Together, these results demonstrate that hippocampal neurons learn the temporal distance structure of the visual sequence, organizing responses smoothly around the reference scene rather than maintaining purely sensory-driven tuning.

**Fig. 2.**
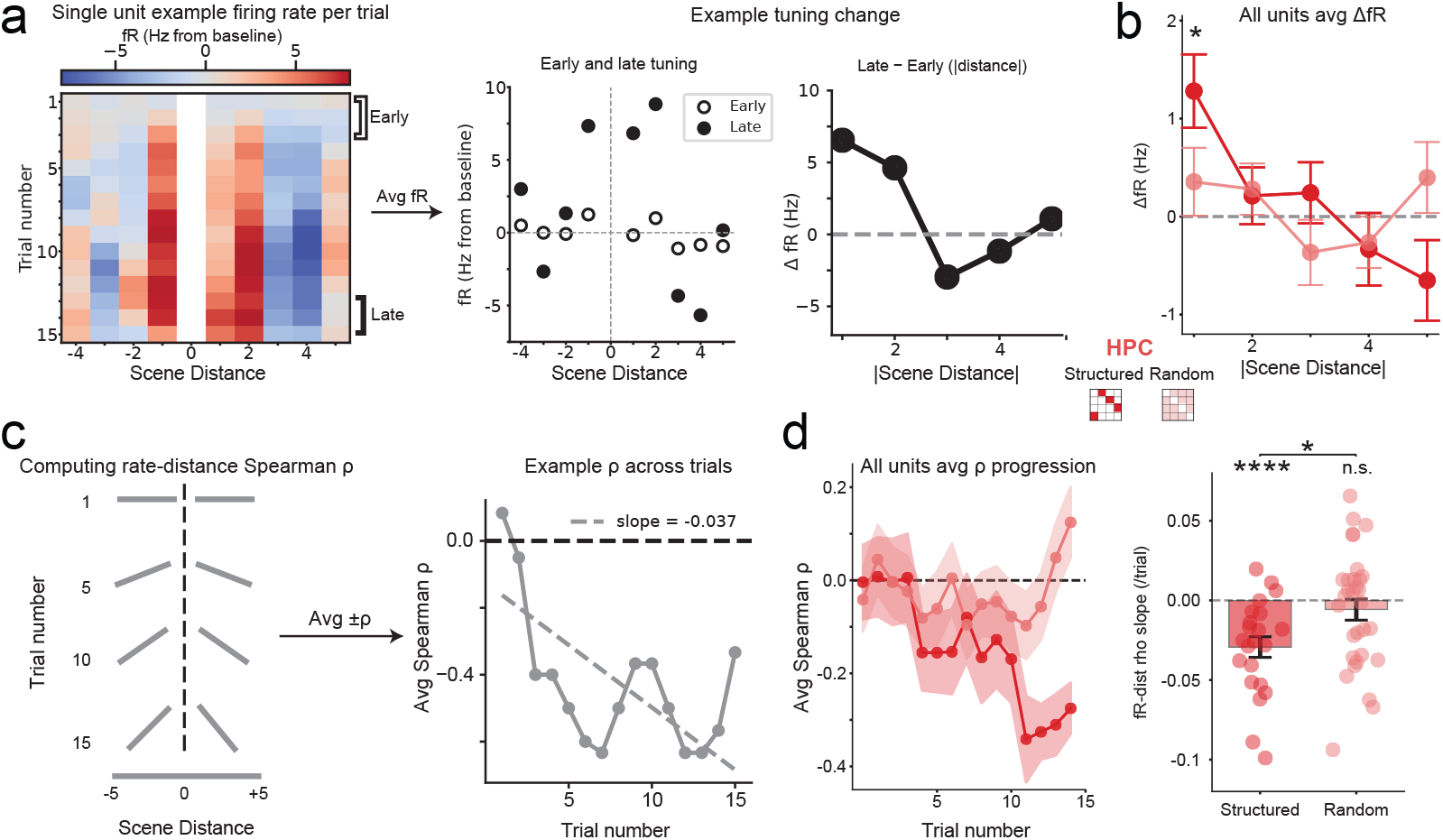
Changes in single-unit activity. (a) Left: Baseline-subtracted firing rates (fR) for an example hippocampal neuron in the structured condition, shown across trials and signed scene distance from the neuron’s reference scene. Distance 0 is omitted to avoid regression-to-the-mean artifacts from peak-based selection. Middle: Early (average of first 3 trials, open circles) and late (average of last 3 trials, closed circles) fR by signed scene distance. Right: Late minus early change in fR as a function of absolute distance. (b) Mean (dots) ± SEM (error bars) of single-unit fR change (late – early) across hippocampal neurons for the structured (red) and random (pink) conditions plotted against absolute distance. (c) Left: For each trial, Spearman’s *ρ* of fR by distance was calculated separately for positive and negative signed distances then averaged. This metric across the 15 trials quantified the emergence of a linearly graded neuronal response from the reference scene. Right: Example hippocampal neuron showing progressively more negative *ρ* across the 15 trials (linear regression line indicated with gray dashes). (d) Left: Mean (dots) ± SEM (bands) of *ρ* between fR and distance across hippocampal neurons for each trial in the structured (red) and random (pink) conditions plotted against trial number. Right: Mean (columns) ± SEM (error bars) of the linear slope between *ρ* and trial number across hippocampal neurons; dots indicate individual neurons in the structured (red) and random (pink) conditions. (**** *p <* 0.0001, * *p <* 0.05, n.s. = not significant)

### Changes in population activity with sequence learning

In addition to changes in single-unit activity, we predicted that visual sequences would be represented at a pop-ulation level in the hippocampus. We defined a vector of responses across hippocampal neurons for each item and repetition and then projected these vectors into a low-dimensional state space to quantify the global geometry of the population code in the hippocampus. The looping of the sequence in the structured condition from end to beginning on each repetition resulted in periodic order of the scenes. Because periodic variables can result in a circular or ring manifold (23–25), we expected that a ring structure would emerge when the hippocampal population vectors for the items in the structured condition were projected into the state space. We used non-linear dimensionality reduction (Isomap) (26) to preserve the intrinsic geometry of the population code and selected the neighborhood size based on when reconstruction error first plateaued (27) (Figure S6).

We used spatial statistics to detect ring structure in state space. A ring corresponds to a scale-specific signature in pairwise distances between population codes: relative clustering at small distances (along its circumference) plus relative dispersion at larger distances (within its hole). Ripley’s *H* function was used to quantify clustering (positive values) and dispersion (negative values) in a 2-D embedding of the population vectors (28) (Figure 3a). Here, *r* refers to the normalized spatial radius in the *H* function, with *r≈* 0 representing the very local distances and *r ≈*1 representing the largest global distances in the given embedding. For each session, we computed *H* curves early (first 3 trials) and late (last 3 trials) in learning and summarized their difference Δ*H* with a scale contrast metric defined as the area under the curve (AUC) at small radii (*r* = 0 to 0.5; reflecting local dispersion) minus AUC at larger radii (*r* = 0.5 to 1; negative values reflecting broader-scale dispersion) (Figure 3b). Positive values of the metric indicate that representations become more clustered locally and more dispersed at broader scales, consistent with spatial rearrangement of the scene manifold towards the shape of a ring.

**Fig. 3.**
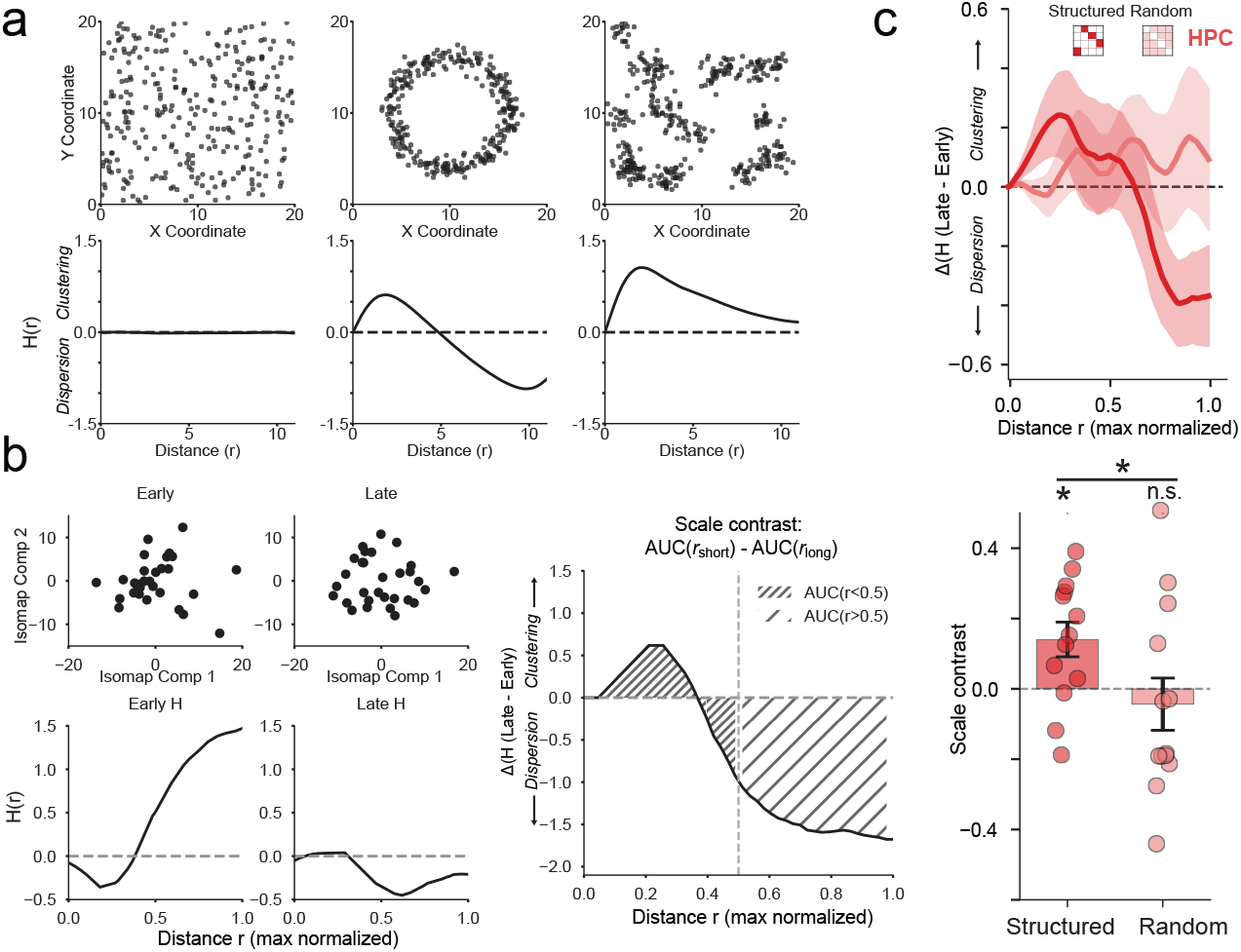
Changes in population activity. (a) Top: Three simulated spatial patterns — Poisson-like uniform (left), noisy ring (middle), and clustered (right) — each represented by 2-D point clouds. Bottom: Ripley’s *H* functions for each simulation; for scale *r, H*(*r*) *>* 0 indicates clustering and *H*(*r*) *<* 0 indicates dispersion. (b) Example hippocampal session in the structured condition. Top: Population vectors of firing rate on early (first 3) and late (last 3) trials projected into a 2-D Isomap component space. Bottom: Empirical Ripley’s *H* curves for early and late phases. Right: Δ*H* (late – early) as a function of normalized distance *r*, with hashed regions marking short-distance (*r*_short_ = 0–0.5) and long-distance (*r*_long_ = 0.5–1) intervals. A scale contrast metric was defined as the difference between the area under the curve (AUC) for short-vs. long-distance Δ*H*. (c) Top: Mean (lines) ± SEM (bands) of Δ*H* curves in the hippocampus across sessions of the structured (red) and random (pink) conditions. Bottom: Mean (columns) ± SEM (error bars) of scale contrast metric across sessions; dots indicate individual sessions in the structured (red) and random (pink) conditions. (* *p <* 0.05; n.s. = not significant)

The hippocampus showed a significant increase in this scale contrast metric for the structured condition (*p* = 0.017; Figure 3c) but not the random condition (*p* = 0.542). Furthermore, the structured condition was significantly greater than the random condition (Mann–Whitney U, *p* = 0.036). No other region showed significant differences between structured and random sequences in evidence for the ring (all *p ≥*0.601; Figure S5).

### Emergence of ring structure in topology of population activity

To examine whether the changes in scale-dependent spatial dispersion reflected a reorganization of the neural manifold, we used persistent homology from topological data analysis to quantify global features of the arrangement of population vectors. Such topological analyses help distinguish true low-dimensional structure from local correlations or sampling noise (29, 30). Persistent homology is a data-driven, non-parametric method that measures the shape of high-dimensional data (here, the population firing-rate vectors corresponding to each scene) using simplicial complexes (31). Simplicial complexes generalize graphs by capturing not only pairwise interactions (edges) but also higher-order relationships (multi-point faces, or simplices) based on geometric criteria such as overlapping radii in the Vietoris–Rips filtration (Figure 4a). By analyzing the topology of these complexes at every possible radius, we can assess connectivity structure across multiple scales. The most characteristic shape of the data is then identified as the topological feature that persists across the greatest range of scales, as evaluated using the persistence diagram (32, 33).

**Fig. 4.**
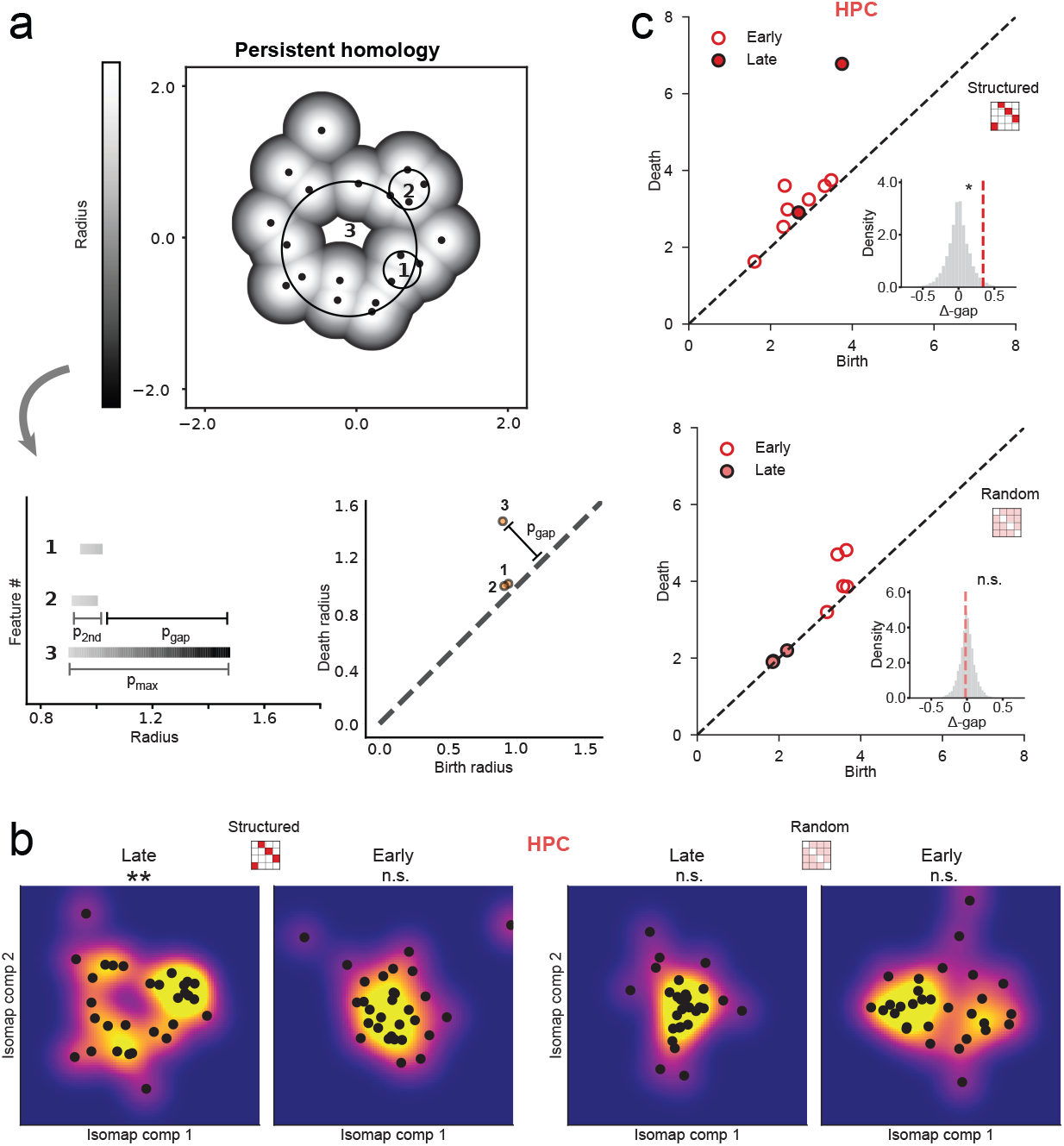
Emergence of topological loop. (a) From the high-dimensional space of population firing-rate vectors (black dots), simplicial complexes form as the connectivity radius (grayscale bar) expands. 1-D topological loops (“features”) appear at a birth radius and disappear at a death radius; these event pairs populate the Betti-1 persistence diagram (bottom). Three hypothetical loops are highlighted with numerals (1–3). The persistence gap (black bar next to feature 2, *p*_gap_) is defined as the difference between the largest (feature 3, *p*_max_) and second-largest (feature 2, *p*_2nd_) persistence values, capturing the relative stability of the dominant cycle. *p*_gap_ was normalized by the midpoint radius of the dominant feature: *r*_dom_ = (*b*_dom_ + *d*_dom_)*/*2 to yield *gap* = *p*_gap_*/r*_dom_. (b) Low-dimensional embedding of hippocampal population activity across all sessions for each of the 10 scenes in the structured condition (left column) and random condition (right column). For each condition, late trials (last 3, left) and early trials (first 3, right) are shown, overlaid with kernel-density estimates (cool = low density, warm = high density). A ring-like manifold emerges only in the structured-late trials, whereas structured-early trials and both phases of the random condition show no significant ring topology (*gap*) compared to a null distribution recomputed from dimension-shuffled population vectors within each phase. (c) Persistence diagrams plot birth versus death radii for early (open) and late (filled) Betti-1 features under structured (top) and random (bottom) conditions. The diagonal marks zero persistence. Insets show null distributions of the Δ-gap (late – early normalized persistence gap) generated by shuffling trial labels (gray, *n* = 50,000), overlaid with the observed Δ-gap (colored dashed line). Significance was computed as the proportion of null values more extreme than the observed value. (** *p <* 0.01, * *p <* 0.05, n.s. = not significant)

We hypothesized that the most characteristic shape of the collection of scene representations late in exposure would be a topological loop (i.e., a circle with a hole) in the hippocampus mirroring the temporally circular sequence in the structured condition. To quantify the prominence of a ring structure, we employed a persistence gap metric, which measures how much the most persistent feature exceeds the next largest persistence (max persistence minus runner-up persistence normalized by the size of dominant feature). Hippocampal population vectors in the structured condition showed a prominent one-dimensional cycle in late exposure (persistence gap compared to dimension-shuffled null distribution; *p* = 0.002; Figure 4b) but not early exposure (*p* = 0.171). No significant cycle was detected in the random condition (early: *p* = 0.771, late: *p* = 0.930). The change in persistence gap with exposure was compared to null distributions generated by randomizing early and late trial labels and recomputing persistence diagrams (Figure 4c). The prominence of topological structure in the hippocampus became significantly larger from early to late trials of the structured condition (Δ*gap, p* = 0.021) but not the random condition (*p* = 0.402). No topological changes were observed in the other recorded brain regions (Figure S7).These findings indicate that the collection of hippocampal scene representations becomes predominated by a stable loop-like topology that mirrors the circular temporal structure of the learned sequence.

### Angular coding of sequence position within ring structure

The topological analyses above demonstrate that the hippocampus represents the sequence in the structured condition in a high-dimensional ring. However, this leaves open the question of where individual scenes fall on the ring. We hypothesized that the ring dimension represents the circular sequence itself and thus that the locations of scenes on the ring are ordered by their serial position. To test this prediction, independent of arbitrary rotation or reflection, we performed a circular–linear regression (34) of scene identity and angular position along the best-fit ellipse within the Isomap space of population vectors (Figure 5a). In the hippocampus, angular position on the ellipse tracked serial position in the sequence (Figure 5b), yielding significantly low regression error (i.e., mean absolute angular deviation between predicted and observed angles) relative to shuffled labels (*p* = 0.004), whereas no such relationship was observed for the random condition (*p* = 0.460). Because non-hippocampal regions did not form a structure ring and thus did not produce interpretable angular positions, we did not apply circular-linear regression to those regions. Together, these analyses show that sequence information is incorporated into hippocampal representations at the level of single-unit responses, the geometry of the global embedding of these patterns, and trajectories along that geometric representation.

**Fig. 5.**
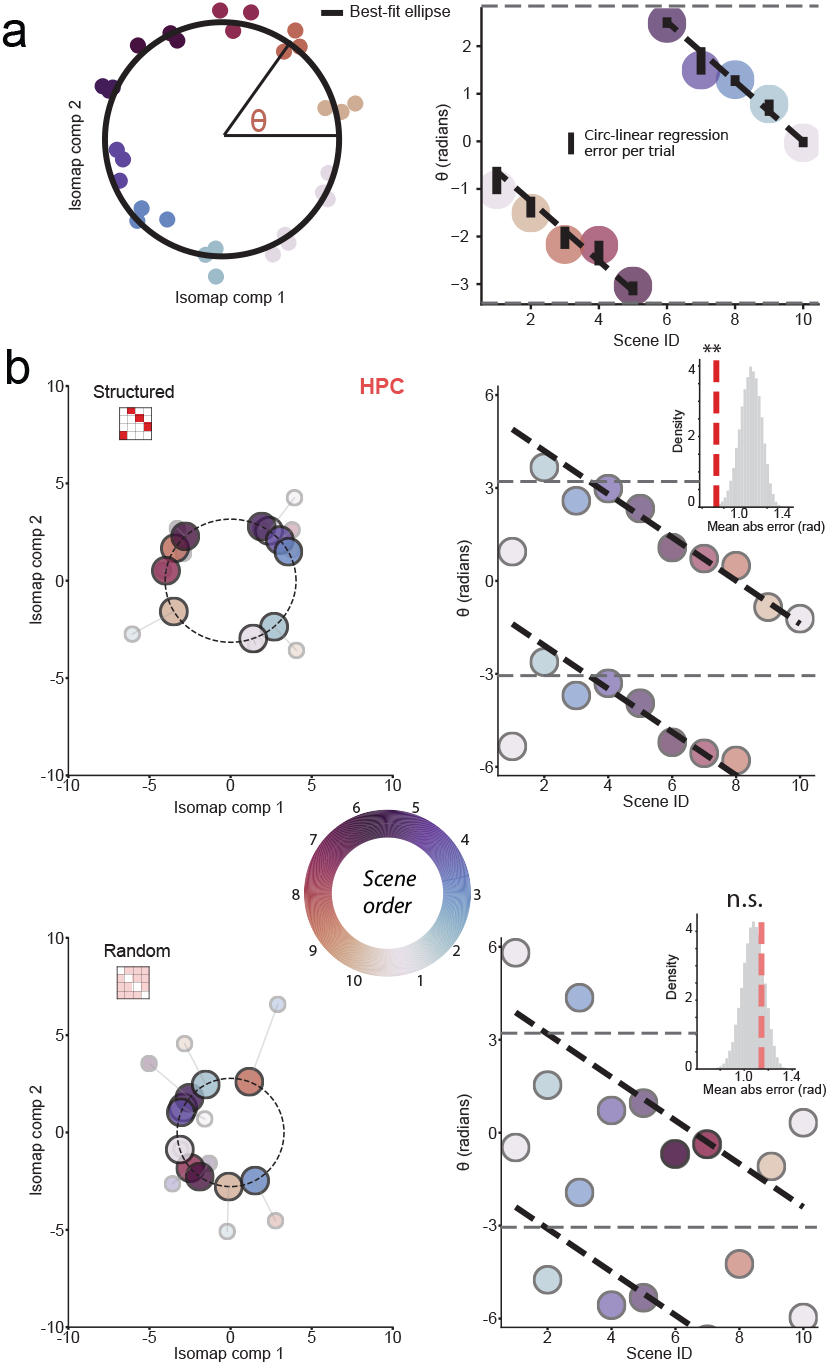
Angular coding of sequence trajectory. (a) An ellipse (black curve) was fitted to the 2-D Isomap embeddings of trial-averaged population vectors (points colored by scene identity). Each point is orthogonally projected onto the ellipse and its angular coordinate *θ* is measured relative to the ellipse’s major axis. These *θ* values serve as circular coordinates in a circular–linear regression relating angle to temporal position in the serial order; absolute residuals (black bars) quantify regression error. (b) Example hippocampal embeddings and regression. Each original point (small, transparent marker) is projected onto the ellipse (large, opaque marker). Top: Isomap embeddings for late trials of structured condition show a clear angular progression consistent with a ring-like trajectory ordered by scene position (1–10), with a well-aligned circular–linear regression (thick dashed line) and low residual error. Gray dashed lines are at *±π*, beyond which half cycles are repeated to visualize circular continuity on a linear axis. Random condition (bottom): The corresponding embeddings lack angular organization and the circular–linear regression shows no consistent relationship between angle and scene position. Insets: Randomization tests of circular–linear regression error in hippocampus, shown separately for structured (top) and random (bottom) conditions during the late trials. For each condition, gray histograms display the null distribution of mean absolute error obtained by shuffling scene labels (*n* = 50,000); dashed vertical red (structured) and pink (random) lines indicate the observed error for with respect to the null distribution and the associated *p*-value. (** *p <* 0.01; n.s. = not significant)

## Discussion

Our findings demonstrate that arbitrary visual sequences can reshape the geometry of sensory codes in human hippocampal neurons. Initially, many hippocampal neurons responded to individual scenes. With repeated exposure to temporally structured sequences, these initial responses evolved such that the neurons responded to other scenes in the sequence in a graded manner inversely proportional to their temporal distance. These changes in tuning did not occur when the sequences were scrambled on each repetition, nor did they occur in the brain regions outside of the hippocampus that we recorded from (amygdala, insula, anterior cingulate cortex). We then used spatial statistics and topological data analysis to reveal a global reorganization of the manifold of hippocampal representations. After exposure to structured but not random sequences, population activity vectors for individual scenes in the hippocampus but not other regions were arranged into a ring that preserved the serial order. These findings suggest that visual sequences are represented in the hippocampus at both single-unit and population levels. Such geometric representations provide a natural substrate for generalization and prediction.

Building on prior fMRI research (19), recent work has demonstrated that hippocampal neurons are sensitive to probabilistic stimulus transitions arising from walks along a graph community structure (20), indicating learning of local temporal regularities. The present results extend this evidence of sensitivity to transition probabilities by showing that hippocampal neurons represent the temporal distance of items in a sequence from an initially preferred reference item. We also added a random control condition that was matched for the types and frequency of scenes but with a scrambled order on every repetition, validating that distance-dependent coding required learning of temporal structure. Moreover, we included single-unit recordings from regions outside of the hippocampus and medial temporal lobe cortex, demonstrating that the effects are anatomically selective to the hippocampus among these regions. Finally, we adopted state-ofthe-art techniques from topological data analysis to provide a new quantitative support for conclusion that the hippocampus embeds temporal structure in the manifold geometry of population activity.

Distance-dependent changes in single-unit activity and the emergence of ring structure in population activity could result from known plasticity mechanisms. According to the non-monotonic plasticity hypothesis (NMPH) (35, 36), the scene that follows a neuron’s reference scene in the structured condition may be integrated into that neuron’s tuning because the response to the reference scene persists into the perception of the neighboring scene, leading to robust coactivation that promotes synaptic strengthening. In contrast, reference scene may be less active during the perception of more distant scenes in the structured sequence, leading to moderate coactivation that promotes synaptic weakening. Behavioral time-scale plasticity (BTSP) (37) and theta sequences (38, 39) may additionally contribute, enabling synaptic changes over behavioral timescales with dendritic plateau potentials and providing temporal compression of event sequences, respectively. The timescale of the learning on the order of minutes is also consistent with rapid extraction of regularities through reactivation within hippocampal pathways (40), which can maintain and reinforce learned structure even within a single session (41, 42). More generally, sequence-dependent changes in tuning and representational geometry could correspond to the development of attractor dynamics in hippocampal circuitry. Recurrent connectivity patterns combining local excitation and longer-range inhibition have been proposed to generate stable activity “bumps” along a temporal dimension (43, 44). Such mechanisms provide plausible explanations for how the hippocampus maintains structured temporal codes that differentiate near from far sequence positions.

Our results also extend the notion of “concept cells” in the hippocampus that can encode abstract, multimodal features (6, 22, 45), by showing that individual cells can also encode sequences of visual stimuli to form flexible memory representations of dynamic environments. Neurons adapt their tuning to reflect temporal relationships, aligning with cortical findings from macaques (15, 17) and with theories that the hippocampus forms conjunctive representations that bind multiple aspects of experience in a context (5, 9–12, 45, 46). By encoding both the identity of stimuli (‘what’) and their temporal order (‘when’), these cells may support episodic memory functions in the hippocampus, which require associating events in sequence.

This work has limitations that highlight important directions for future research. Recordings were obtained from patients with epilepsy who had clinically determined electrode placement, resulting in uneven sampling across medial temporal lobe regions and limiting circuit-level or causal inferences (47, 48). Although we excluded data from within 4 hours of a seizure, our results may not generalize to the non-epileptic brain and may be infuenced by prescribed medications. Moreover, this study involved short sequences from one sensory modality (vision) that were repeated in quick succession and contained deterministic probabilities. It will be important to assess the generality of our findings with longer, spaced, and multimodal sequences that more closely resemble natural environments. Finally, we used a passive viewing task to allow implicit sequence learning and to simplify the procedure for the patient population. As a result, we did not have any behavioral measures of attention or learning to corroborate the neural effects. We inferred sequence learning because the neural effects emerged over time and were stronger than in a control condition without temporal structure. Subjects were continuously monitored by research personnel during testing and encouraged (when needed) to remain engaged and minimize distraction. However, directly relating neural and behavioral measures of sequence learning in future work will be helpful for establishing downstream functional relevance (49).

Together, these findings demonstrate that visual sequences are learned rapidly in the human hippocampus by encodeing the temporal distance between items in single-unit activity and by representing the geometrical structure of sequences in the state space of population activity. By showing that relationships among neuronal responses come to mirror the temporal relationships among sequence elements, this work provides a concrete representational framework by which hippocampal activity can form memories that unfold over time. Disruption of such representations may contribute to the hippocampal-dependent temporal-order and relational memory deficits observed in Alzheimer’s disease and related conditions (50–52).

## Methods

### Study participants

Seventeen patients (7 males and 10 females, 16 right-handed and 1 left-handed, ages 15 to 61 years) with medication-refractory epilepsy undergoing intracranial EEG electrode implantation for seizure localization were enrolled in the study after providing informed consent (see Table S2 for subject demographics). Electrode planning and implantation was made exclusively for clinical purposes. Behnke-Fried depth electrodes (Ad-Tech Medical Instrument Corp., Oak Creek, WI, USA) with 40 uM microwires extending from the tip were used to record single units from the hippocampus (HPC), amygdala (AMY), anterior insula (AIC), and anterior cingulate cortex (ACC) (Figure 1a, Table S1, S2). All participants remained on their home antiseizure medications for the duration of the study. The study was approved by the Institutional Review Board at Yale University.

### Task design

Participants were exposed to sequences of visual scenes while neuronal activity was recorded. Visual stimuli consisted of 20 photographs of natural scenes, based on the importance of the human hippocampus for scene perception (53). Two types of sequences were constructed: in the structured condition, 10 of the scenes were selected to form a fixed sequence that repeated in the same order 15 times, looping from the end to the beginning; in the random condition, the other 10 scenes were presented in each of the 15 repetitions, but their order was randomized on each cycle to eliminate temporal structure. Each scene was displayed for 1 s, followed by a fixation cross for 1 s. The order of the conditions was counterbalanced across participants.

### Electrophysiological recording

Voltage traces were high-pass filtered at 300 Hz for spike detection. Spikes were then sorted offline with Waveclus (54). Sorting parameters were calibrated per participant to account for variability in siginal amplitude and background noise. Putative single units were retained if they showed stable waveforms and had fewer than 5% of spikes with inter-spike intervals under 3 ms. We recorded a total of 134 single units from the hippocampus, 89 units from amygdala, 108 units from insula, and 113 units from ACC. Spike-sorting quality metrics confirmed well-isolated units across regions (Figure S1). Each neuron was recorded during both conditions (structured and random) in all but two sessions in which only one condition could be obtained. Because not all neurons were recorded and responsive in both conditions, we compare conditions across partially overlapping sets of neurons rather than within individual units.

### Defining scene-responsive reference units

We aligned spike times to scene onset (time 0) and calculated firing rates within a 1000-ms post-stimulus window, subtracting a 1000-ms pre-stimulus baseline (fixation cross period). To assess learning-related changes at the single-unit level, we first selected units that met a minimum average firing rate threshold of 1 Hz during at least one scene across trials. We then defined a reference scene for each unit as the scene in that condition that elicited the highest mean firing rate early in sequence exposure (first 3 trials). A unit was classified as responsive to its reference scene if: (i) its maximum firing rate evoked by the scene was more than 3 standard deviations greater than the fixation baseline (after 50 ms standard deviation Gaussian kernel smoothing was applied separately to baseline and response periods), (ii) its mean firing rate evoked by the scene was greater than the 95th percentile of a null distribution generated by shuffling scene labels 50,000 times, and (iii) the firing rate evoked by the preceding (-1) and following (+1) neighbors was no more than 5% of the response to the reference scene. These inclusion criteria were designed to be less stringent than traditional definitions of visual selectivity across many trials (22, 55), reflecting our goal of defining a reference point for learning-related changes in a limited number of early trials rather than stable stimulus selectivity.

For each early scene-responsive unit, firing rates in early (first three) versus late trials (last three) were compared across structured and random conditions to assess how sequence learning modulated tuning. Importantly, we focused on changes in firing rate to scenes in the sequence other than the reference soon, to avoid regression to the mean or other selection biases, and because our hypothesis concerned the impact of sequence learning on increased tuning to neighboring scenes. Moreover, the selection procedure was identical for the structured and random conditions, so any statistical artifacts from this procedure cannot explain differences between conditions. Moreover, the population analyses below include all recorded units regardless of early responsiveness further validating that the overall claims do not depend on the inclusion criteria.

Analyses of single-unit scene responsiveness based on mutual information (MI) were conducted separately for hippocampus, amygdala, insula, and dorsal anterior cingulate cortex (Figure S3a). MI was computed using `mutual_info_score` from the scikit-learn library between scene labels and spike counts during scene exposure periods. Statistical significance was assessed with a permutation test in which scene labels were randomly shuffled within trial, yielding a null distribution of surrogate MI values (50,000 iterations); p-values were calculated as the proportion of surrogate MI values exceeding the true MI value. Units were classified as visually responsive at *α <* 0.05 for comparison with prior literature.

To assess the stability of early scene responsiveness across the full session, MI was additionally compared between the first 3 trials (early MI) and all trials (session MI). MI values were z-scored within unit relative to corresponding scene label shuffled null distributions (50,000 iterations). Normalized early MI was binned (in steps of 0.2 z) and the mean corresponding session MI was calculated for each bin. The relationship between early and session MI was evaluated using Pearson correlation and least-squares regression separately for each brain region (Figure S3b).

### Distance dependence of single-unit firing rates

For each early-scene responsive unit and each of the 15 trials, we quantified distance-dependent tuning using an averaged monotonicity index. This index was calculated as the average of two Spearman rank correlations (*ρ*): the correlation across scenes ahead of the reference scene in the sequence between their firing rate and their (positive) serial order distance (+1 to +5), and the negation of the correlation across scenes preceding the reference scene in the sequence between their firing rate and their (negative) scene distance (-1 to - 4). Trial-wise estimates were computed after smoothing firing rates with a three-trial sliding window, and the average *ρ* was Fisher z-transformed to enable comparisons across trials. This construction yields a signed, within-trial measure of whether responses decrease monotonically with increasing absolute distance from the reference scene, independent of overall response magnitude.

To assess learning-related changes, we summarized the temporal evolution of the monotonicity index for each unit by fitting a linear regression as a function of trial number, yielding a single learning slope per unit. Statistical significance of learning-related changes was evaluated using Wilcoxon signed-rank tests, and differences between structured and random conditions were assessed using Mann–Whitney U tests.

### Spatial statistics of representations (Ripley’s H)

For each session, population activity patterns were constructed by concatenating the firing rates of all simultaneously recorded units within a given brain region and condition for each trial and scene. These scene-specific population vectors were projected into a two-dimensional (2-D) space using Isomap with *k* = 7 nearest neighbors. The parameter *k* was chosen by finding the smallest neighborhood size at which reconstruction error plateaus, defined as when two consecutive values of *k* first show less than 5% improvement in error (Figure S6). Beyond this plateau, added neighbors no longer contribute independent distance information, implying that intrinsic local connectivity has been captured.

To quantify the spatial organization of these embeddings, Ripley’s H function — a normalized measure of point-pattern clustering at each distance scale — was computed separately for the population maps from early (first 3 trials) and late (last 3 trials) phases of sequence exposure. Distances were normalized within each session so that the maximal inter-point distance equaled 1. The difference between late and early H curves (Δ*H* = *H*_*late*_*− H*_*early*_) was smoothed across distance bins to obtain session-level Δ*H* profiles. For each Δ*H* profile, an excursion metric was defined as the signed area difference between short-range (*r* = 0 to 0.5) and long-range (*r* = 0.5 to 1.0) portions of the curve, computed via numerical integration. Positive values indicate greater increase of local clustering relative to long-range dispersion. Excursion metrics were computed for every session and averaged within condition by brain region. Between-condition differences were evaluated using two-sided Mann–Whitney U tests.

### Topological data analysis (TDA) of scene representations

TDA is a nascent research field, drawing upon concepts from computational topology to describe the “shape” of data at multiple scales in terms of persistent homology. Given a dataset, persistent homology first calculates a topological approximation of the data. Such an approximation typically takes the form of a simplicial complex, which is a data structure that generalizes a graph beyond dyadic interactions by incorporating higher-order relations (simplices) between three or more elements. Here, we used a Vietoris– Rips complex, which is a simplicial complex consisting of progressively more fine-grained representations of a dataset. This construction is followed by the calculation of topological shape features like connected components, cycles, and voids, which are ultimately represented in a compact 2-D representation the persistence diagram. A point (*x, y*) in a persistence diagram denotes a topological feature being “born” at scale *x* and “destroyed” at scale *y*. Despite its low dimensionality, this representation is expressive and provably stable to noise in the data. Moreover, persistence diagrams permit filtering the original data based on the idea of discerning between signal and noise, which was shown to be particularly relevant in the context of data analysis (33).

We performed persistent homology analyses on population firing-rate representations of scenes across trials (as used above) to assess the emergence of a ring-like structure in hippocampal activity (Figure 4). Vietoris–Rips complexes were constructed using Ripser (56), and persistence diagrams were computed to characterize topological features across scales. 1-D holes were identified via Betti-1 features. To quantify the prominence of the dominant loop relative to both noise and overall scale, we computed a normalized persistence gap. For each persistence diagram, Betti-1 features were ranked by lifetime (death *−* birth). If no features were detected the persistence gap was 0. Otherwise, the persistence gap (*p*_*gap*_) was defined as the difference between the largest and second-largest lifetimes (32, 33), normalized by the midpoint radius of the dominant feature *r*_dom_ = (*b*_dom_ + *d*_dom_)*/*2 to yield *gap* = *p*_*gap*_*/r*_dom_. This normalization ensures that loop stability is evaluated relative to the scale of the dominant feature in the embedding. Null distributions of *gap* were generated by shuffling the dimensions of the population vectors (50,000 iterations). To isolate learning-related changes in loop stability, we computed the change in the normalized persistence gap from the early to the late phase, Δ*gap* = *gap*_late_ *−gap*_early_. Null distributions of Δ*gap* were generated by shuffling the phase label (50,000 iterations). Observed values were compared against these null distributions to obtain two-sided *p*-values.

### Ordering of scene identities on topological ring

Population firing-rate representations from the late phase for each scene were embedded into Isomap component space using the same procedure as above. The best fit ellipse of the resulting point cloud in each condition was computed. Circular linear regression (34) of the angle of each point on the ellipse with the serial position of that scene was computed. To ensure that the regression captured appropriately spaced progression around the ring rather than trivial solutions involving multiple cycles, regression slopes were restricted to values consistent with a single traversal, corresponding to approximately one scene step per 2*π*/10 radians, allowing at most *±*1 scene deviation over a full cycle. The fit of the regression was quantified as the mean absolute angular deviation between predicted and observed angles. Errors were computed per trial and averaged across trials. Statistical significance was assessed with permutation tests in which scene labels were shuffled within trials (50,000 iterations). *p*-values were computed as the proportion of iterations with corresponding error less than the observed error.

## Data and code availability

Analyses were conducted using Python (version 3.9.16) with NumPy, Pandas, SciPy, Matplotlib, and scikit-learn libraries. The data from this study are freely available without stipulation or restrictions by email request to the corresponding author. Original code is available at https://github.com/damisahlab/Hippocampal-Sequences. Any additional information required to reanalyze the data is available from the corresponding author upon request.

## Author contributions

Conceptualization, E.C.D., and N.B.T-B.; methodology, A.A., T.B., E.C.D., and N.B.T-B.; investigation, T.B., A.A., Y.Z., B.R., N.B.T-B., and E.C.D.; writing-–original draft, T.B.; writing-–review & editing, T.B., E.C.D., B.R., and N.B.T-B.; funding acquisition, E.C.D.; resources, E.C.D., and N.B.T-B.; supervision, B.R., E.C.D., and N.B.T-B.

## ACKNOWLEDGEMENTS

We thank the Yale Comprehensive Epilepsy Center clinical team and the patients who participated. This work was funded by a Hypothesis Fund grant (E.C.D.), National Institute of Health grants R01MH138291 (E.C.D.) and R01MH069456 (N.B.T-B.), the Canadian Institute for Advanced Research (N.B.T-B.). and the Wellcome Trust Grant: 310142/Z/24/Z (ECD).

## Supplementary Information

**Fig. S1.**
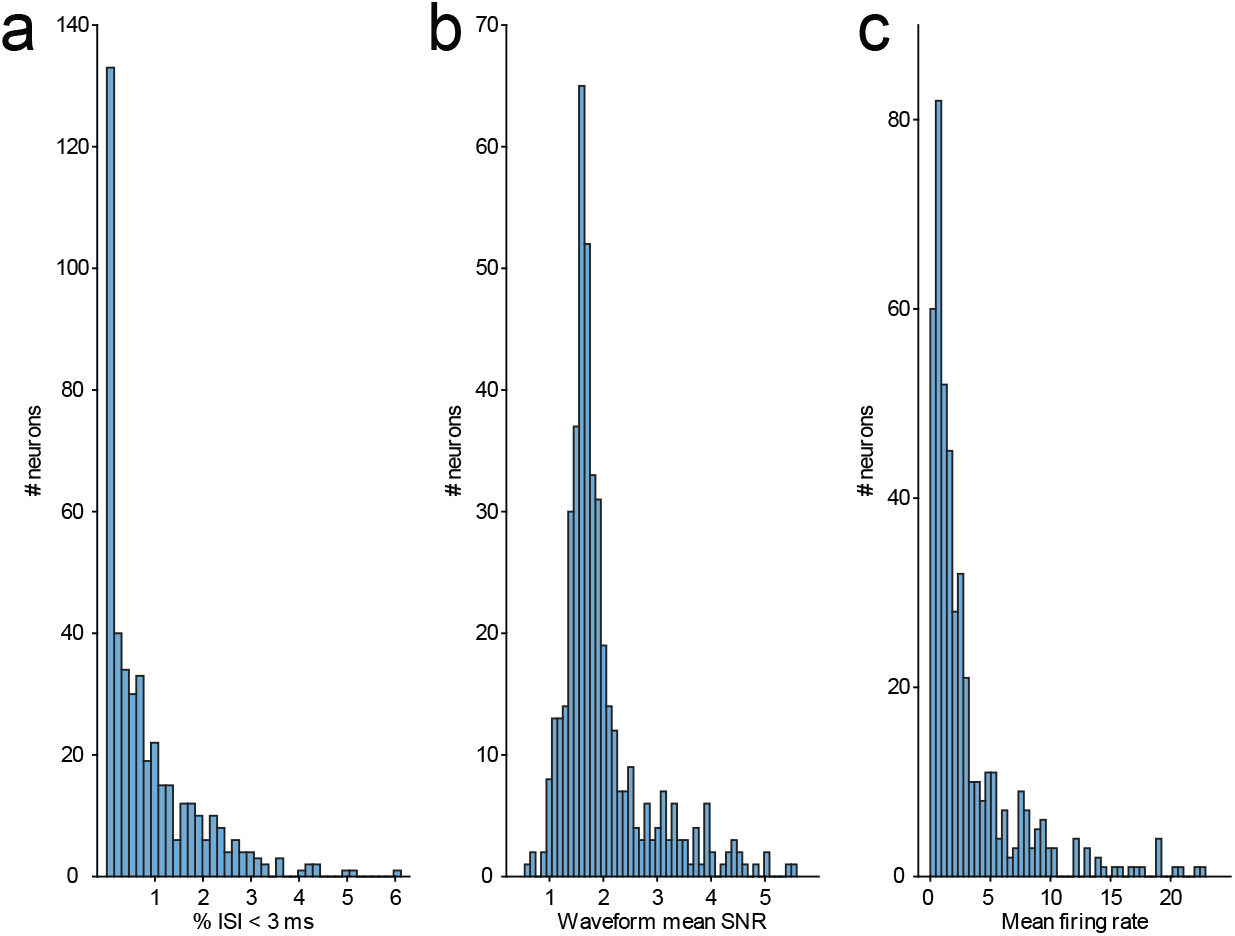
Spike sorting quality metrics for all detected single units. (a) Histogram of inter-spike intervals (ISI) below 3 ms (mean = 0.85%, s.d. = 0.99%). (b) Histogram of signal-to-noise ratio (SNR) between mean amplitude of waveform and s.d. of noise (1.97 *±* 0.81). (c) Histogram of mean firing rate across the entire recording session (3.17 *±* 3.97).

**Fig. S2.**
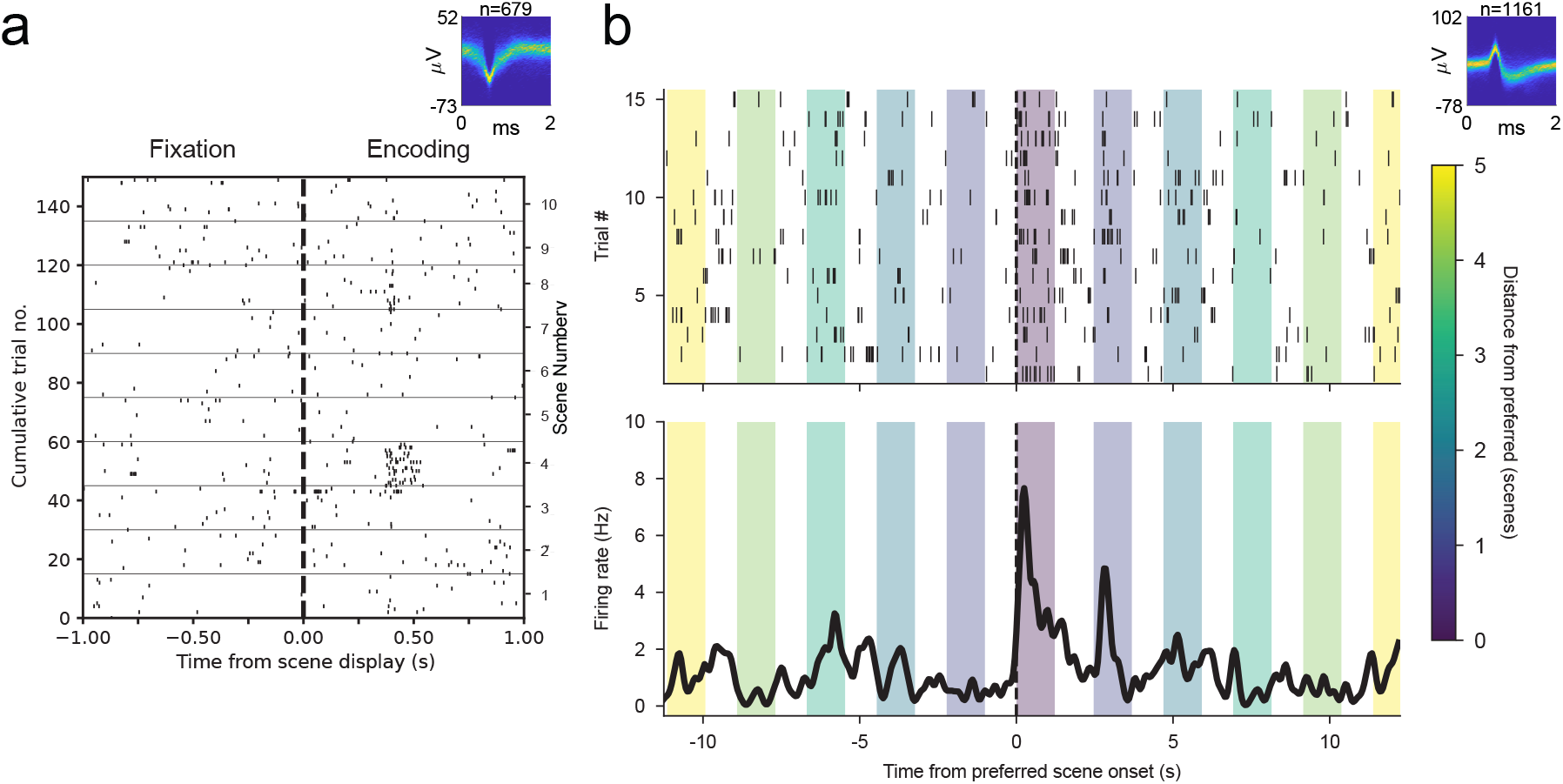
Example single-unit responses in random and structured conditions. (a) Example hippocampal neuron recorded during the random condition. Inset depicts voltage waveforms over time for all spikes from this unit. Raster plot of this unit’s within-trial spiking activity aligned to scene onset (time 0; dashed line). Each row corresponds to one trial (left axis) and trials are grouped by scene identity (right axis); the 15 trials for each scene are ordered sequentially by repetition from bottom to top. (b) Example hippocampal neuron recorded during the structured condition. Inset depicts voltage waveforms over time for all spikes from this unit. Top: Raster plot aligned to the onset of the neuron’s initially responsive reference scene (time 0; dashed line). Each row corresponds to one of the 15 sequence repetitions, with trials ordered from bottom to top (left axis). The color of background shading indicates the distance of neighboring scenes from the reference scene. Bottom: Peristimulus time histograms (PSTHs) of firing rate over sequence averaged across all trials.

**Fig. S3.**
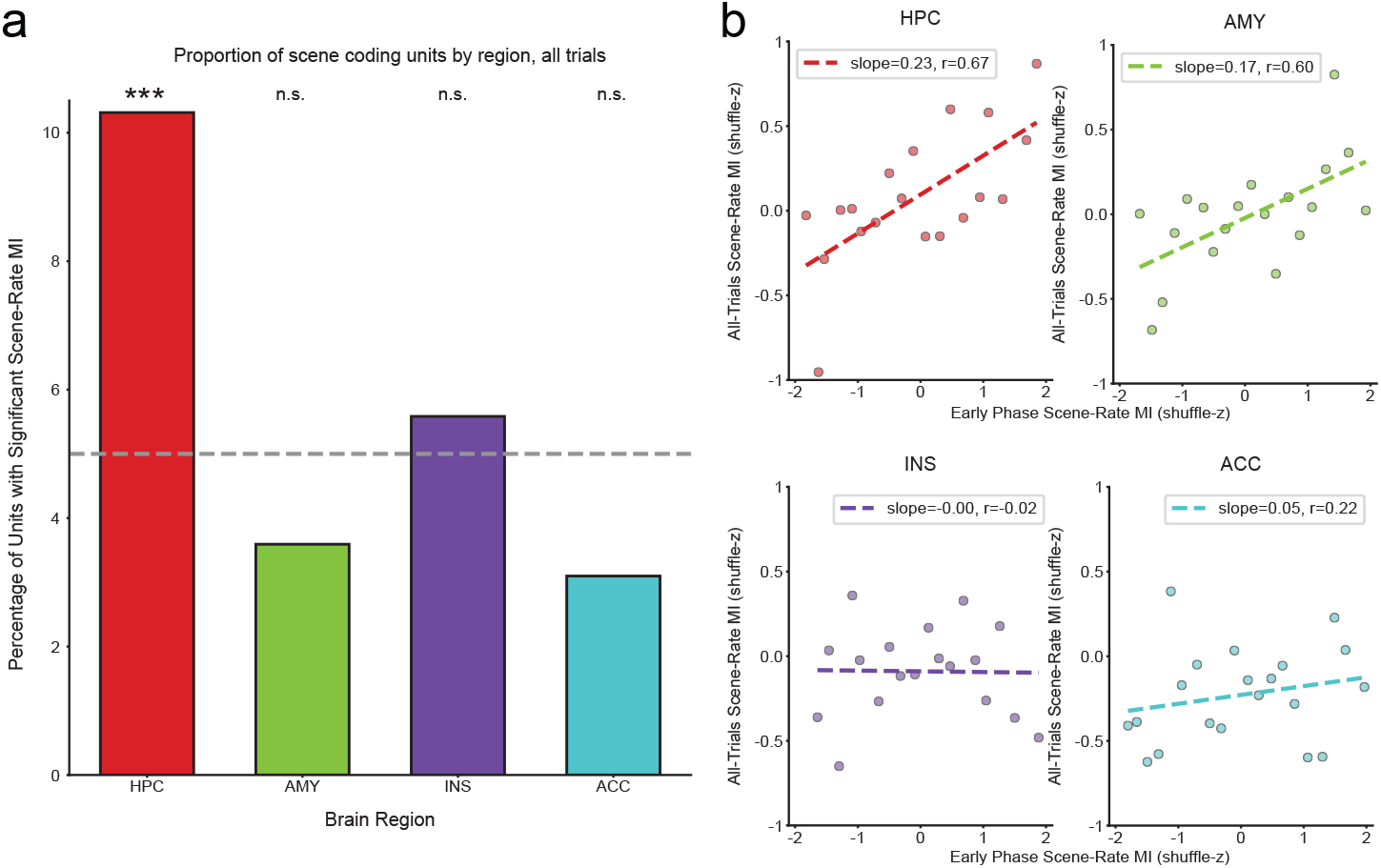
Validation of visual scene selectivity proportions using mutual information (MI). (a) Proportion of single units exhibiting significant MI between firing rate and scene identity, computed across all trials, shown separately for hippocampus (HPC), amygdala (AMY), insula (INS), and anterior cingulate cortex (ACC). MI was computed between integer-valued scene-evoked response matrices (10 scenes by 15 trials) and scene identity, and statistical significance was assessed using a randomization test based on shuffled scene labels (50,000 iterations; *α* = 0.05, dashed line). The percentage of units with significant MI differed across brain regions (*χ*^2^ = 12.88, *p* = 0.0049). Only the hippocampus showed a significantly higher percentage of scene-responsive neurons than expected by chance (HPC: +3.64*σ*; AMY: *−*0.83*σ*; INS: +0.38*σ*; ACC: *−*1.31*σ*), where *σ* is the standard deviation of the null distribution computed as the normal approximation to a binomial model with underlying *p* = 0.05. (b) Early MI was calculated from the first three presentations of each scene (z-scored within each unit relative to a label shuffled null distribution of 50,000 iterations), the same trials used to define reference scenes based on initial responsiveness in the main single-unit analysis. Units were grouped into bins of 0.2 z and the mean early MI and overall MI (all trials) values are plotted with a linear fit (dashed line). The robust positive correlation in the hippocampus indicates that the initial responsiveness of units is relatively stable throughout the session. (*** *p <* 0.001; n.s. = not significant)

**Fig. S4.**
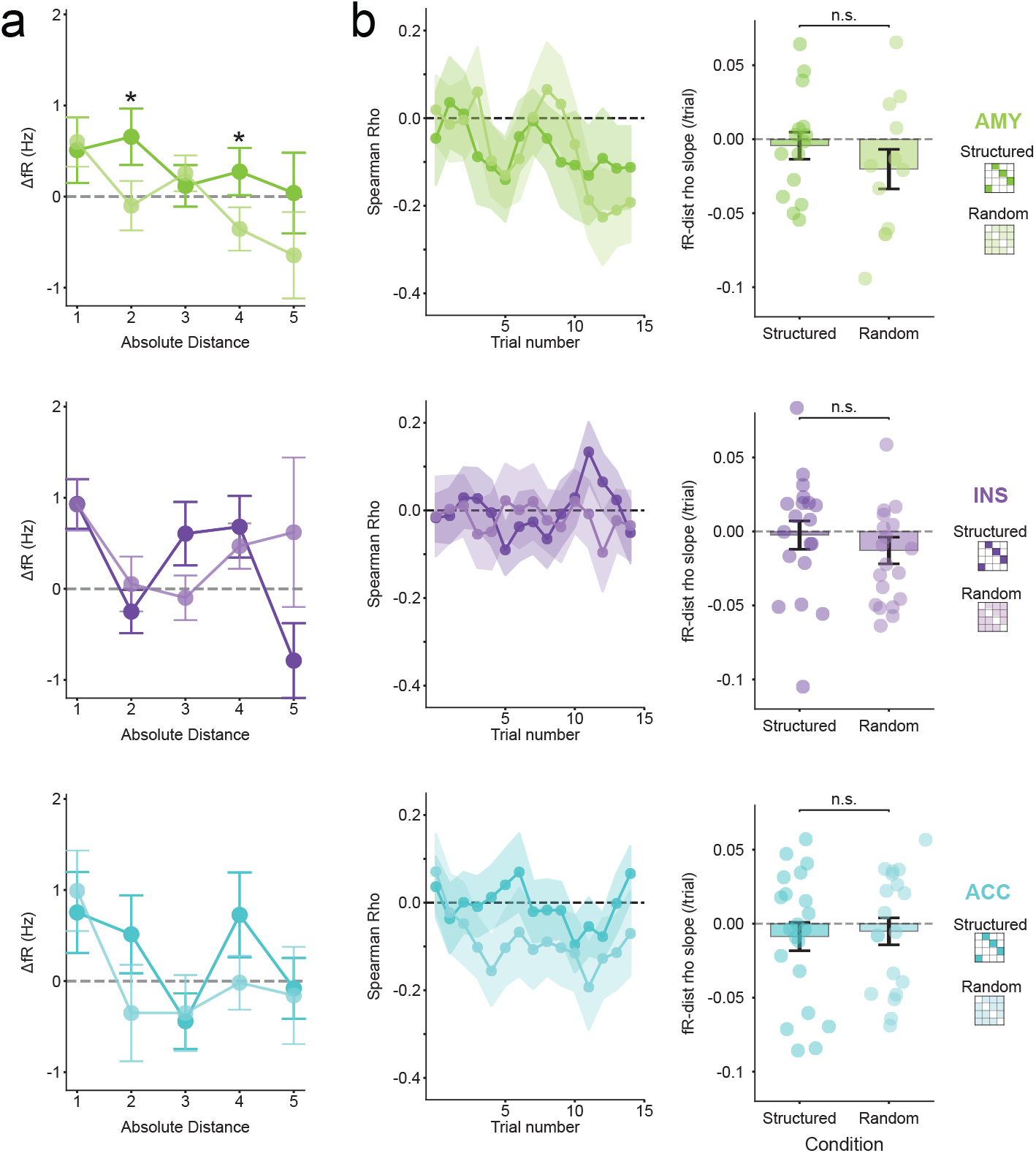
Analyses from Figure 2 in non-hippocampal regions. (a) Average late–early change in firing rate by absolute scene distance for amygdala (AMY, green), insula (INS, purple), and anterior cingulate cortex (ACC, cyan) in the structured (dark) vs. random (light) conditions. No non-hippocampal region showed a reliable increase in response to the neighboring +1 scene nor a monotonic decrease with distance, effects that were observed in the hippocampus; AMY showed greater change for structured vs. random at +2 and +4 distances, suggesting a more global increase rather than distance coding. (b) Accordingly, no non-hippocampal region showed a reliable change in distance-dependent coding across trials for structured vs. random conditions, as observed in the hippocampus. (* *p <* 0.05; n.s. = not significant)

**Fig. S5.**
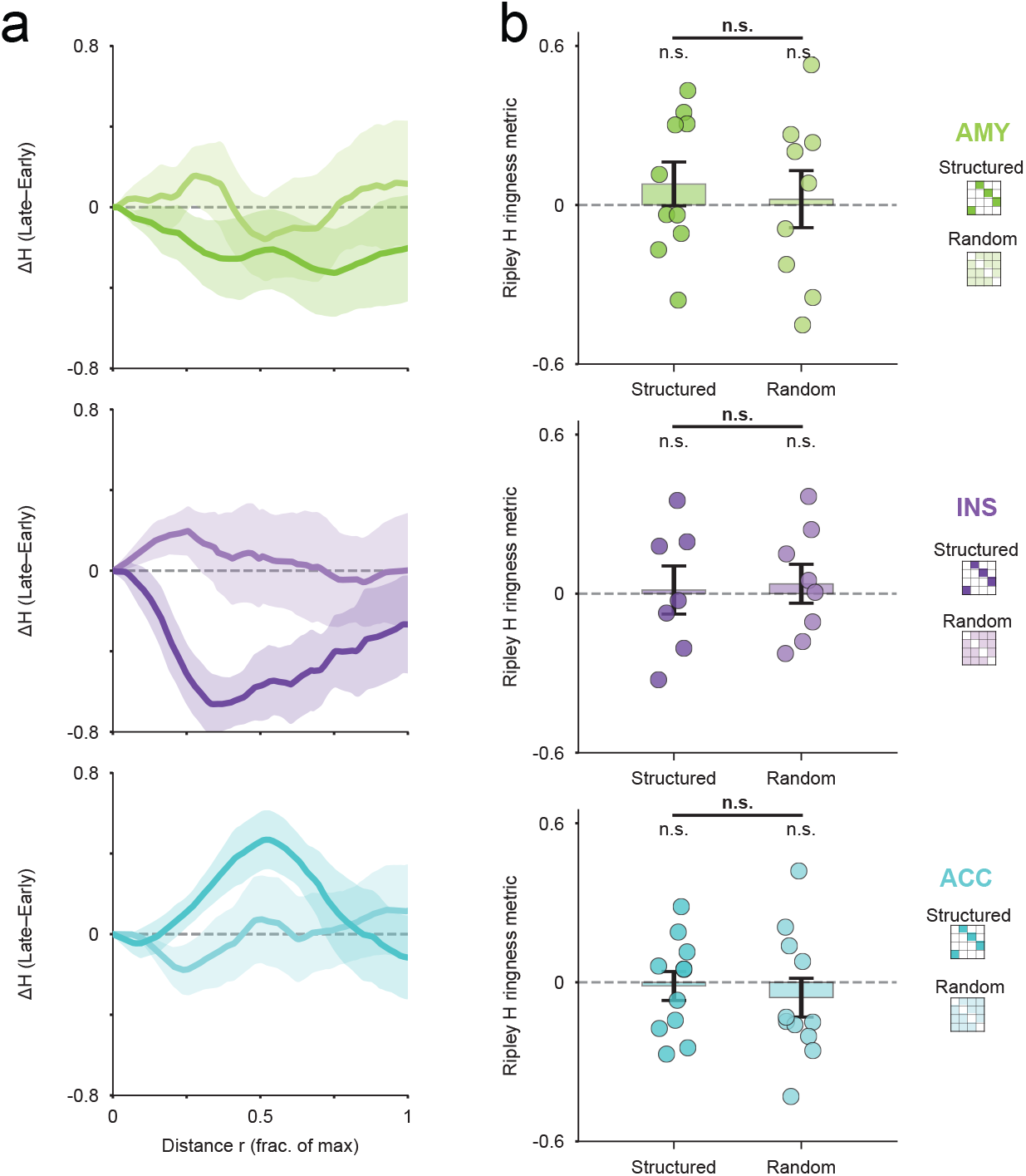
Analyses from Figure 3 in non-hippocampal regions. (a) Change in Ripley’s H (late – early) as a function of normalized distance (fraction of maximum *r*) for population codes in the amygdala (AMY, green), insula (INS, purple), and anterior cingulate cortex (ACC, cyan). Curves show mean ± SEM across sessions for structured (dark) and random (light) conditions. Unlike HPC, non-HPC regions did not exhibit a systematic short-distance increase plus long-distance decrease in H indicative of a ring-like geometry. (b) Session-level scale contrast metric (difference between short-vs long-distance area under the Δ*H* curve) for structured and random conditions. Bars denote mean ± SEM; points indicate individual sessions. (n.s. = not significant)

**Fig. S6.**
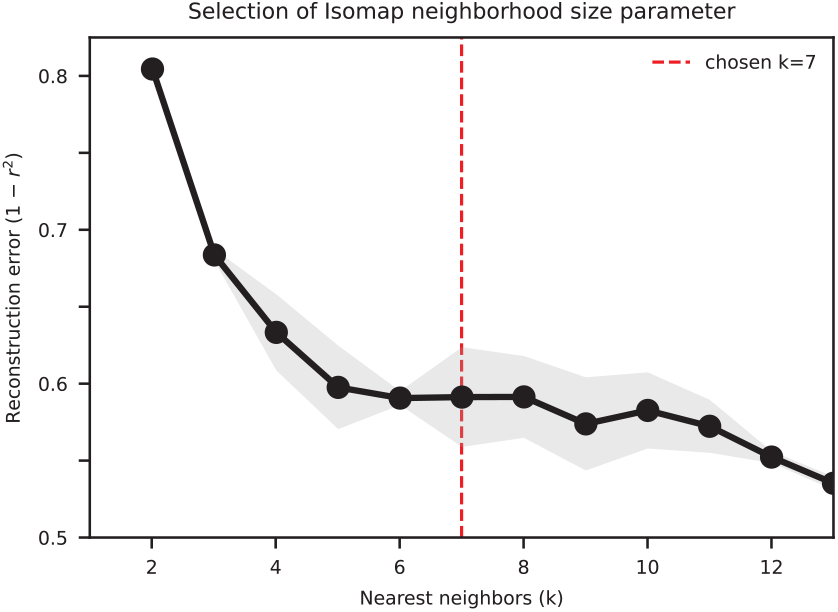
Selection of Isomap neighborhood size. Neural population activity from the hippocampus in both conditions and for both early and late trials was embedded using Isomap, and the neighborhood size *k* was selected based on reconstruction error. For each condition, firing-rate feature matrices were reduced with PCA (retaining the number of components whose mean cumulative variance explained was closest to 67 percent across conditions), and Isomap embeddings were computed over a range of nearest-neighbor values (*k* = 2 to 12). Reconstruction error was quantified as 1 *− r*^2^, where *r* denotes the correlation between pairwise distances in the original high-dimensional space and in the Isomap embedding. Curves show the reconstruction error averaged across conditions, with shading indicating ± SEM. To identify an appropriate neighborhood size, we evaluated the relative improvement in reconstruction error as *k* increased and defined a plateau as a reduction of less than 5% for two consecutive values of *k*. The chosen neighborhood size (red dashed line) corresponds to this plateau, yielding a conservative embedding that preserves global structure without over-smoothing.

**Fig. S7.**
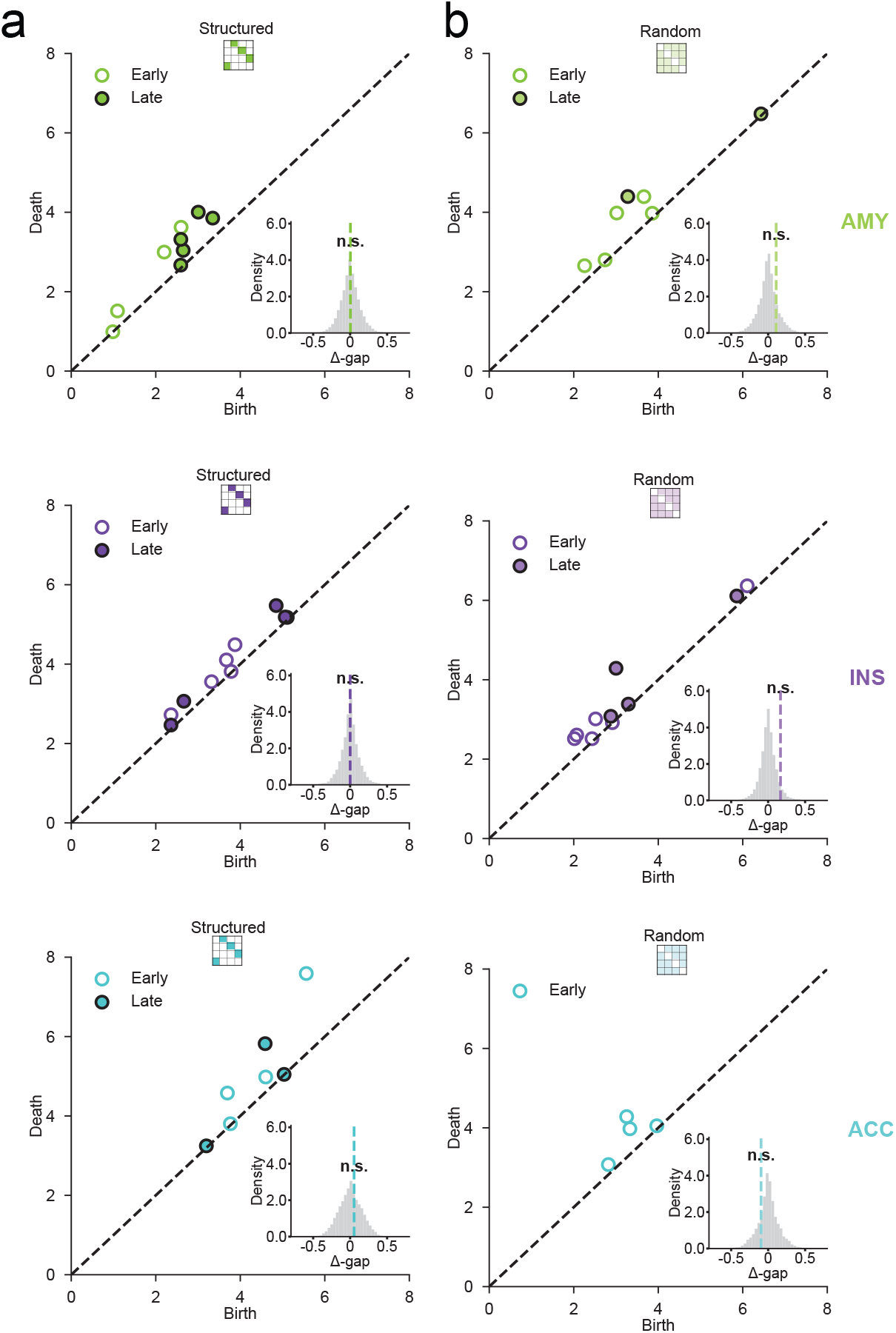
Analyses from Figure 4 in non-hippocampal regions. Persistence diagrams (birth vs. death radii) for Betti-1 features computed from population firing-rate vectors in amygdala (AMY), insula (INS), and anterior cingulate cortex (ACC), shown separately for structured (left column) and random (right column) conditions. Early (open symbols) and late (filled symbols) trial features are plotted for each session where detected; the diagonal denotes zero persistence. Insets show null distributions of the change in normalized persistence gap (Δ*gap*); the true value for each region and condition is indicated with a colored dashed line. No region showed a significant increase from early to late trials in persistence gap for the structured condition. (n.s. = not significant)

**Table S1.** MNI coordinates of microwires for different brain regions.

| Electrode ID | Subject ID | Location | Side | X | Y | Z |
| --- | --- | --- | --- | --- | --- | --- |
| 1 | 1 | Hippocampus | Left | -25.68 | -18.76 | -19.12 |
| 2 | 1 | Amygdala | Left | -20.63 | -5.92 | -20.86 |
| 3 | 1 | Insula | Left | -34.58 | 11.45 | -8.61 |
| 4 | 1 | Insula | Left | -35.13 | 3.54 | -5.99 |
| 5 | 2 | Hippocampus | Left | -26.11 | -9.44 | -23.71 |
| 6 | 2 | Hippocampus | Right | 24.24 | -10.36 | -19.10 |
| 7 | 2 | Insula | Left | -36.68 | 6.32 | -6.98 |
| 8 | 2 | Anterior Cingulate | Left | -7.13 | 25.32 | 19.77 |
| 9 | 2 | Anterior Cingulate | Right | 6.44 | 25.40 | 21.98 |
| 10 | 3 | Hippocampus | Right | 25.41 | -12.53 | -17.62 |
| 11 | 3 | Hippocampus | Right | 31.39 | -28.20 | -6.98 |
| 12 | 3 | Hippocampus | Left | -28.83 | -21.17 | -13.09 |
| 13 | 3 | Amygdala | Right | 20.59 | -1.82 | -24.19 |
| 14 | 3 | Amygdala | Left | -20.49 | -5.15 | -21.68 |
| 15 | 3 | Anterior Cingulate | Right | 4.55 | 27.17 | 23.09 |
| 16 | 4 | Hippocampus | Left | -23.11 | -14.84 | -15.39 |
| 17 | 4 | Hippocampus | Left | -22.35 | -31.47 | -4.94 |
| 18 | 4 | Hippocampus | Right | 26.51 | -15.82 | -15.76 |
| 19 | 4 | Amygdala | Left | -16.84 | -5.02 | -18.27 |
| 20 | 4 | Insula | Left | -37.77 | 6.53 | -15.26 |
| 21 | 4 | Anterior Cingulate | Left | -3.58 | 22.47 | 28.05 |
| 22 | 5 | Hippocampus | Left | -19.09 | -7.05 | -25.35 |
| 23 | 5 | Hippocampus | Right | 23.33 | -12.54 | -20.00 |
| 24 | 5 | Amygdala | Left | -19.86 | 1.47 | -27.02 |
| 25 | 5 | Amygdala | Right | 19.99 | 0.05 | -28.69 |
| 26 | 5 | Insula | Right | -32.76 | 3.22 | 1.80 |
| 27 | 5 | Insula | Right | 42.79 | 14.86 | -4.98 |
| 28 | 5 | Insula | Right | 40.55 | 3.19 | -0.37 |
| 29 | 5 | Anterior Cingulate | Left | -4.53 | 21.13 | 25.53 |
| 30 | 5 | Anterior Cingulate | Right | 11.29 | 25.12 | 27.94 |
| 31 | 6 | Hippocampus | Left | -24.66 | -18.37 | -17.27 |
| 32 | 6 | Hippocampus | Right | 27.60 | -16.62 | -17.10 |
| 33 | 6 | Amygdala | Left | -18.97 | -6.17 | -20.79 |
| 34 | 6 | Amygdala | Right | 24.96 | -6.31 | -25.30 |
| 35 | 6 | Insula | Left | -29.55 | 8.10 | -2.65 |
| 36 | 6 | Insula | Left | -35.32 | 7.10 | -14.52 |
| 37 | 6 | Insula | Right | 33.57 | 5.57 | -3.40 |
| 38 | 6 | Insula | Right | 44.70 | 9.50 | -10.58 |
| 39 | 7 | Hippocampus | Right | 37.76 | -13.55 | -18.30 |
| 40 | 7 | Amygdala | Right | 16.08 | -2.12 | -22.97 |
| 41 | 7 | Anterior Cingulate | Left | 0.68 | 30.83 | 26.48 |
| 42 | 8 | Hippocampus | Right | 32.02 | -25.76 | -5.58 |
| 43 | 8 | Insula | Right | 35.43 | 12.44 | 2.62 |
| 44 | 9 | Hippocampus | Left | -21.00 | -16.91 | -19.73 |
| 45 | 9 | Amygdala | Left | -15.93 | -5.97 | -24.35 |
| 46 | 9 | Anterior Cingulate | Left | -2.61 | 27.77 | 15.63 |
| 47 | 9 | Anterior Cingulate | Right | 7.62 | 28.86 | 15.03 |
| 48 | 10 | Amygdala | Left | -19.35 | -5.77 | -21.51 |
| 49 | 10 | Hippocampus | Left | -24.07 | -20.87 | -20.33 |
| 50 | 10 | Hippocampus | Right | 30.58 | -20.10 | -18.43 |
| 51 | 10 | Anterior Cingulate | Left | -39.08 | 10.70 | -11.28 |
| 52 | 10 | Anterior Cingulate | Right | 40.10 | 8.92 | -8.68 |
| 53 | 11 | Amygdala | Right | 15.91 | -4.56 | -23.79 |
| 54 | 11 | Hippocampus | Left | -23.46 | -19.53 | -15.36 |
| 55 | 11 | Hippocampus | Right | 29.22 | -20.59 | -13.60 |
| 56 | 11 | Anterior Cingulate | Left | -2.51 | 29.88 | 20.64 |
| 57 | 11 | Anterior Cingulate | Right | 4.24 | 30.10 | 21.77 |
| 58 | 11 | Insula | Left | 35.07 | 7.50 | -3.56 |
| 59 | 12 | Amygdala | Right | 19.27 | -3.14 | -18.30 |
| 60 | 12 | Hippocampus | Right | 25.22 | -9.19 | -20.95 |
| 61 | 12 | Anterior Cingulate | Right | 5.77 | 30.51 | 27.34 |
| 62 | 12 | Insula | Right | 44.67 | 17.21 | -10.85 |
| 63 | 13 | Anterior Cingulate | Left | -7.64 | 27.89 | 20.93 |
| 64 | 13 | Insula | Left | -35.73 | 19.06 | -7.16 |
| 65 | 13 | Hippocampus | Left | -33.66 | -33.05 | -8.26 |
| 66 | 13 | Hippocampus | Left | -31.56 | -14.61 | -16.81 |
| 67 | 14 | Amygdala | Right | 19.80 | -1.10 | -20.07 |
| 68 | 14 | Hippocampus | Right | 33.51 | -14.97 | -17.82 |
| 69 | 15 | Anterior Cingulate | Left | -4.17 | 34.54 | 14.62 |
| 70 | 15 | Hippocampus | Left | -24.78 | -15.64 | -20.24 |
| 71 | 16 | Amygdala | Left | -17.35 | -4.73 | -19.21 |
| 72 | 16 | Hippocampus | Left | -20.95 | -10.75 | -24.31 |
| 73 | 16 | Anterior Cingulate | Left | -2.57 | 24.57 | 18.89 |
| 74 | 16 | Insula | Left | -33.26 | 4.51 | -4.16 |
| 75 | 17 | Amygdala | Right | 16.89 | -1.96 | -23.04 |
| 76 | 17 | Hippocampus | Right | 26.17 | -17.73 | -18.69 |

**Table S2.** Subject information and detected neuron count per region.

| Subject ID | Age | Handedness | Sex | Total units | HPC | AMY | ACC | INS |
| --- | --- | --- | --- | --- | --- | --- | --- | --- |
| 1 | 23 | R | Male | 29 | 25 | 2 | 0 | 2 |
| 2 | 36 | R | Male | 40 | 20 | 0 | 13 | 7 |
| 3 | 35 | L | Female | 33 | 8 | 12 | 13 | 0 |
| 4 | 45 | R | Female | 36 | 5 | 1 | 29 | 1 |
| 5 | 27 | R | Female | 43 | 19 | 13 | 3 | 8 |
| 6 | 61 | R | Female | 26 | 16 | 5 | 0 | 5 |
| 7 | 28 | R | Male | 11 | 2 | 3 | 6 | 0 |
| 8 | 15 | R | Female | 3 | 2 | 0 | 0 | 1 |
| 9 | 41 | R | Female | 22 | 6 | 9 | 7 | 0 |
| 10 | 23 | R | Male | 41 | 6 | 16 | 0 | 19 |
| 11 | 31 | R | Female | 91 | 5 | 15 | 31 | 40 |
| 12 | 32 | R | Male | 11 | 11 | 0 | 0 | 0 |
| 13 | 22 | R | Male | 9 | 1 | 0 | 0 | 8 |
| 14 | 23 | R | Female | 13 | 4 | 8 | 1 | 0 |
| 15 | 44 | R | Female | 2 | 0 | 0 | 2 | 0 |
| 16 | 39 | R | Female | 14 | 0 | 1 | 8 | 5 |
| 17 | 38 | R | Male | 20 | 4 | 4 | 0 | 12 |

